# MYC is essential for induction of major ZGA and subsequent preimplantation development

**DOI:** 10.1101/2023.06.06.543968

**Authors:** Takuto Yamamoto, Haoxue Wang, Hana Sato, Shinnosuke Honda, Shuntaro Ikeda, Naojiro Minami

## Abstract

In mouse preimplantation development, zygotic genome activation (ZGA), which synthesizes new transcripts from the embryos, begins in the S phase of the one-cell stage, with major ZGA occurring especially at the late two-cell stage. *Myc* is a transcription factor expressed in parallel with ZGA, but its direct association with the major ZGA has not been clarified. In this study, we found that developmental arrest occurs at the two-cell stage when mouse embryos were treated with antisense oligos targeting *Myc* or inhibitors specific for MYC from the one-cell stage. In order to identify when MYC inhibition affected development, we applied time-limited inhibitor treatment, and found that inhibition of MYC at the two-cell, four-cell, and morula stages had no effect on preimplantation development, whereas treatment with the inhibitor at the early two-cell stage arrested development at the two-cell stage. Furthermore, transcriptome analysis revealed that when MYC function was inhibited, genes expressed in the major ZGA phase were suppressed. These results suggest that Myc is essential for the induction of major ZGA and its subsequent development. Revealing the function of *Myc* in preimplantation development is expected to contribute to advances in assisted reproductive technology.

## Introduction

The *Myc* family of transcription factors consists of *Myc, Mycn* and *Mycl*. In most cases, these components form a heterodimer with MAX (Myc-associated factor X) via the leucine zipper domain and bind to target E-boxes for transcription elongation^1,2^. In mammals, newly fertilized embryos use mRNA and proteins stored in the cytoplasm of the oocytes to sustain subsequent development^3–5^. In the mouse embryo, these mRNAs and proteins are degraded as development progresses, and two rounds of embryonic genome activation (ZGA), called minor ZGA and major ZGA, initiate the synthesis of new transcripts from the embryo. The former occurs during the S phase of the one-cell stage, and the latter occurs late in the two-cell stage^6–9^. Immunofluorescence staining analysis confirmed that MYC protein was detected in the nuclear speckle, which controls pre-mRNA splicing immediately after transcription, and was localized in the nucleus in growing oocyte (GO) and fully grown oocyte (FGO), as well as in fertilized eggs (embryos) up to the morula stage, but the signal intensity was slightly weakened at the morula stage and no signal was observed at the blastocyst stage^10^. Only a few studies have attempted to determine the function of *Myc* in preimplantation development by inhibition, but these studies have produced inconsistent results due to different timing of inhibition and different molecular targets.

It has been reported that when *Myc* mRNA was degraded by antisense oligos (ASOs) from the two-cell stage, developmental arrest occurs at the eight-cell/morula stage^11^. Homozygous mutations in *Myc* have been shown to be lethal between embryonic days 9.5 and 10.5^12^. It has also been reported that inhibition of MYC-MAX dimerization by inhibitors of MYC from 2 hours after in vitro fertilization (IVF) resulted in suppression of minor ZGA and developmental arrest mainly at the one-cell stage^13^. However, the risk of nonspecific effects should always be considered when using inhibitors. These reports indicate that embryonic development can proceed to E9.5 with maternally expressed *Myc* alone, and that inhibition of both maternal and embryonic *Myc* results in developmental arrest during preimplantation development. However, since the previous study inhibited MYC from the two-cell stage, when ZGA has already begun, and since the analysis used only inhibitors, the possibility that the phenotype was an artifact could not be ruled out, and thus the exact role of MYC in preimplantation development could not be analyzed.

In this study, we treated one-cell stage (3hpi) embryos with *Myc*-targeted ASOs or inhibitors and investigated their subsequent development. Furthermore, after determining when ASOs and inhibitors affect embryonic development, we also examined the effect of MYC inhibition on gene expression.

## Results

### Knockdown of Myc mRNA in one-cell embryos results in embryonic lethality at the two-cell stage

The expression pattern of *Myc* mRNA in mouse MII oocytes and preimplantation embryos was examined. RT-qPCR analysis of *Myc* mRNA showed that expression levels increased from 6 h post-insemination (hpi) to 48 hpi (Fig. 1A), consistent with a previous report analyzing published RNA-seq data^14^. To investigate the role of *Myc* in early embryonic development, *Myc* was knocked down in mouse preimplantation embryos. *Myc*-targeted antisense oligos (ASOs: *Myc*-ASO-1 and *Myc*-ASO-2) were microinjected into embryos at 3 hpi and cultured until 96 hpi. RT-qPCR confirmed a significant decrease in *Myc* mRNA expression at 24 hpi (Fig. 2B), and immunofluorescence staining showed that MYC was not incorporated into the nuclei of *Myc* knocked down embryos at 24 hpi (Fig. 2C). Morphological observation showed that the developmental rate of *Myc*-ASOs microinjected embryo to the four-cell stage was markedly reduced compared to the control (98.3% for non-targeted control (NT)-ASO, 7.9% for *Myc*-ASO-1 and 1.6% for *Myc*-ASO-2 at 48 hpi) (Fig. 1D and 1E).

**Fig. 1.**
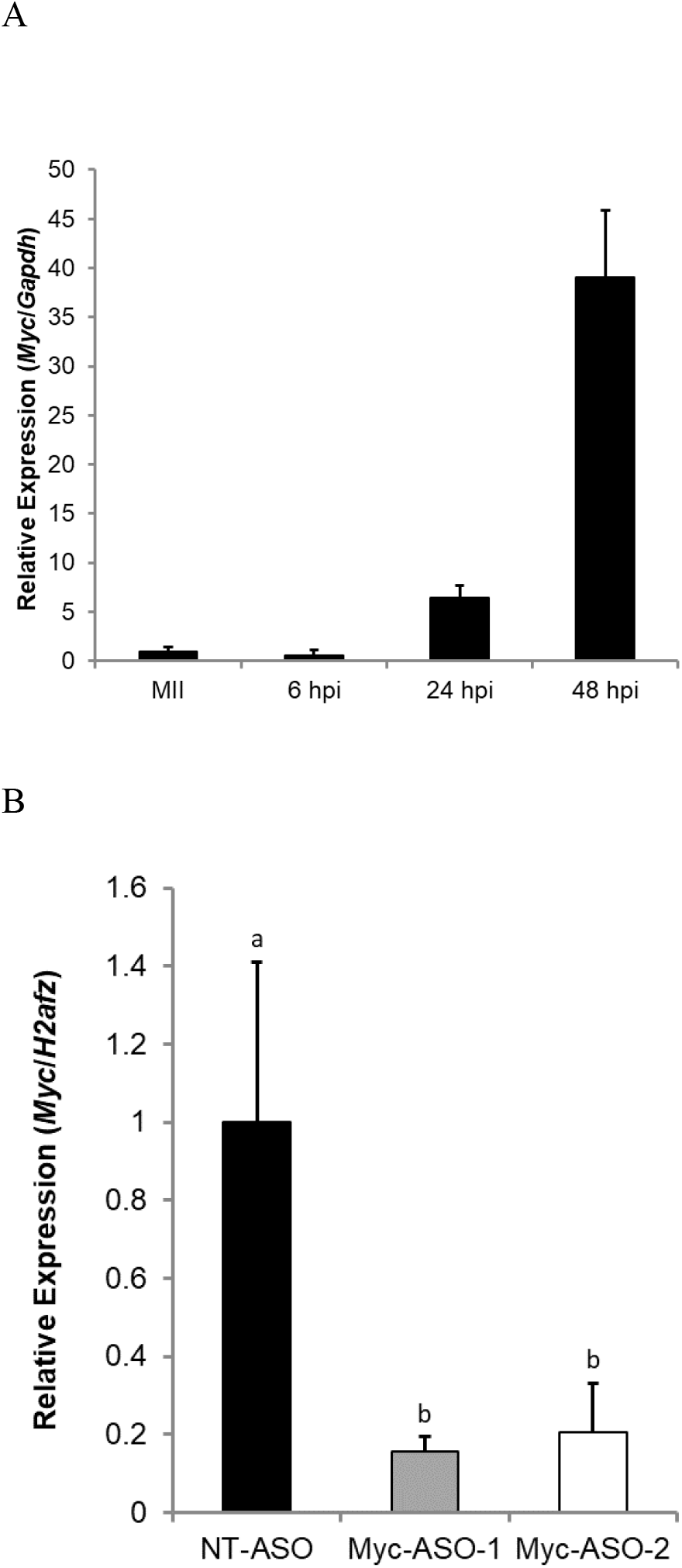
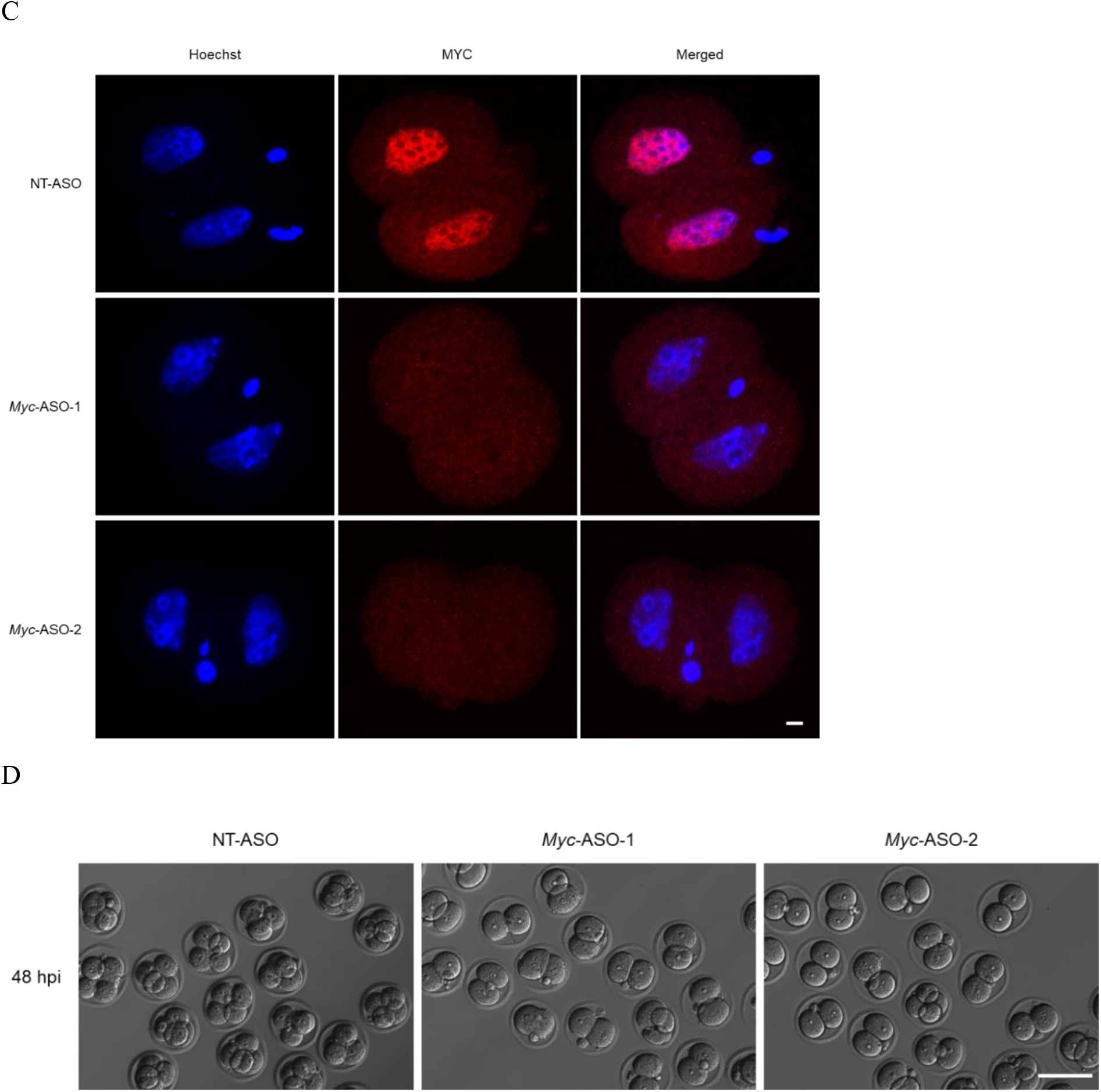
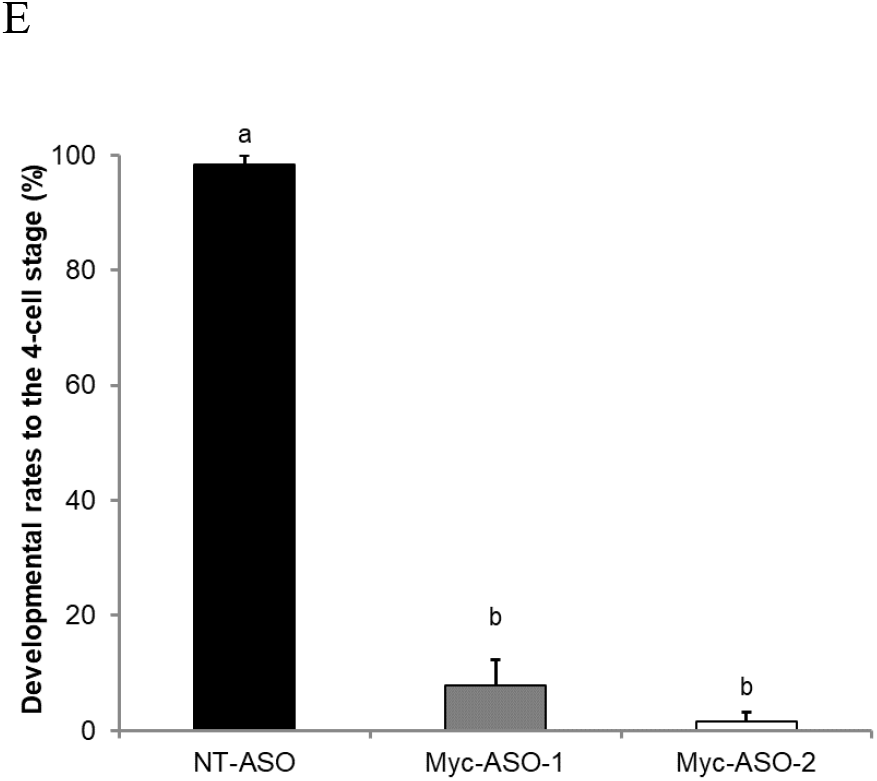
Effects of inhibiting *Myc* mRNA expression on preimplantation development. (A) Reverse transcription-quantitative PCR (RT-qPCR) analysis of *Myc* mRNA expression in metaphase II (MII) oocytes and preimplantation embryos. Gene expression levels were normalized to *Gapdh* as an internal control. Data are expressed as means ± standard error of the mean (S.E.M) (N=3). Thirty oocytes or embryos were analyzed in each treatment. (B) RT-qPCR analysis of *Myc* mRNA expression in *Myc*-knockdown embryos and control embryos at 24 h post-insemination (hpi). Gene expression levels were normalized to *H2afz* as an internal control. Data are expressed as means±S.E.M (N=3). Fifteen embryos were analyzed in each treatment. Statistical analysis was performed using one-way analysis of variance (ANOVA) followed by the Tukey-Kramer test. P values < 0.05 were considered statistically significant and are represented by different letters (a and b). (C) Localization of MYC in embryos injected with either *Myc*-targeted antisense oligos (*Myc*-ASO)-1 (n=16), *Myc*-ASO-2 (n=15) or non-targeted control (NT-ASO) (n=12) at the one-cell stage. MYC is localized in the nuclei of embryos injected with NT-ASO. However, MYC localization in the nuclei disappeared in embryos injected with *Myc*-ASOs. Embryos were photographed at 24 hpi. Scale bar, 20 μm. (D) Representative images of embryos injected with either *Myc*-ASO-1, *Myc*-ASO-2 or NT-ASO at the one-cell stage. Embryos were photographed at 48 hpi. Scale bar, 100 μm. (E) Developmental rates of embryos injected with either *Myc*-ASO-1 (n=63), *Myc*-ASO-2 (n=62) or NT-ASO (n=58) at the one-cellstage. Dataare expressed as means ±S.E.M (N=3). Statistical analysis was performed using the chi-squared test with Holm’s adjustment. P values < 0.05 were considered statistically significant and are represented by different letters (a and b).

**Fig. 2.**
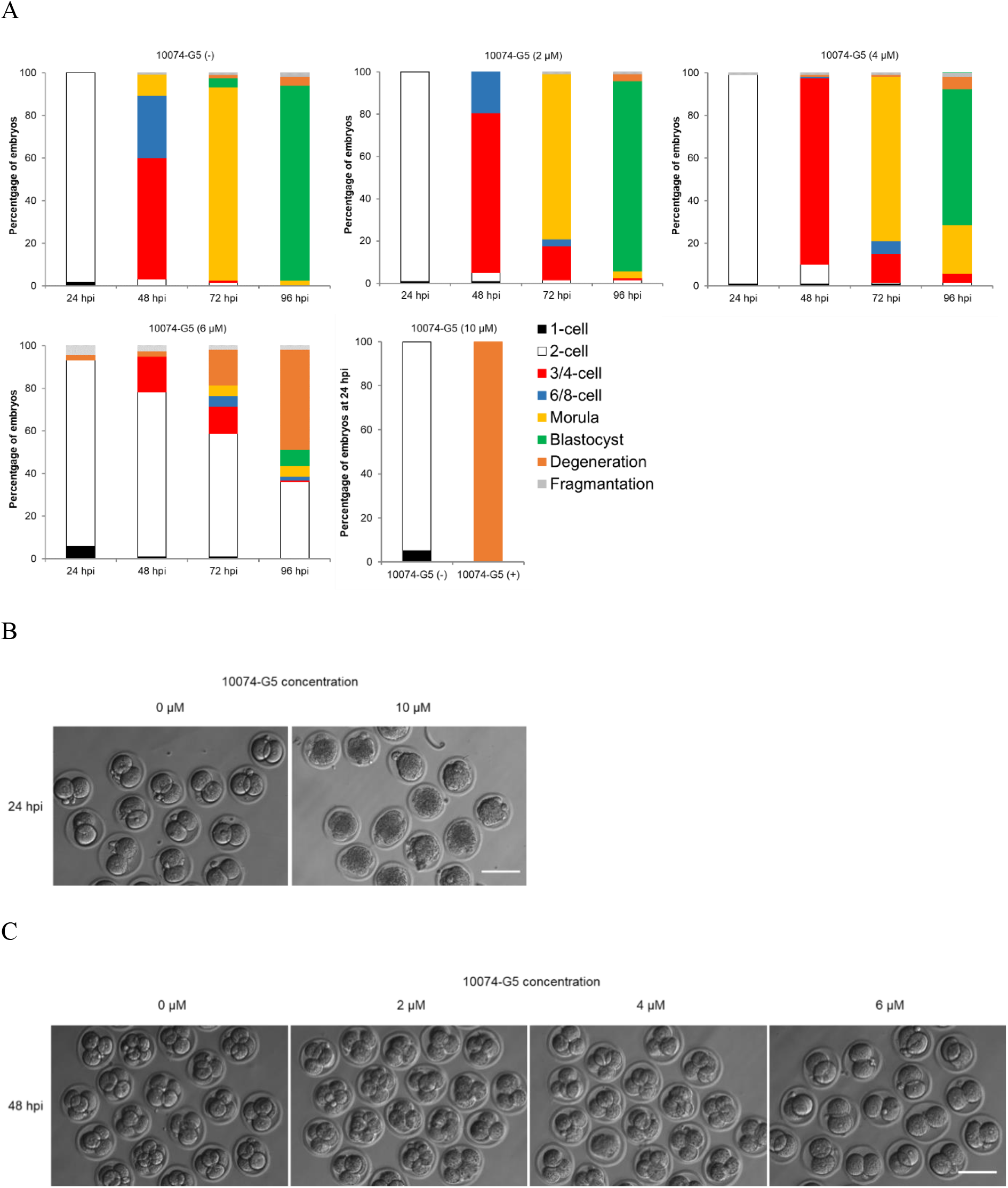
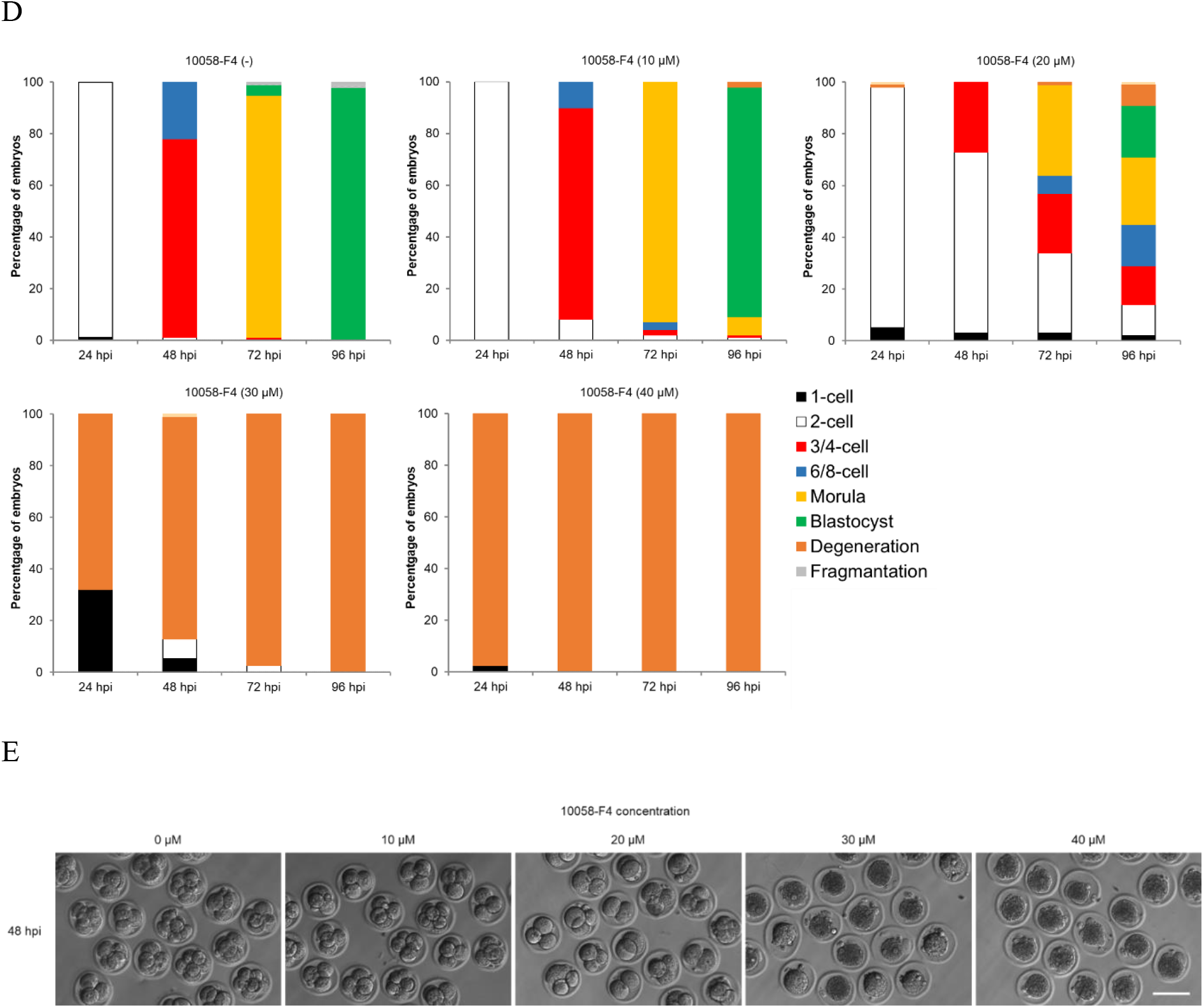
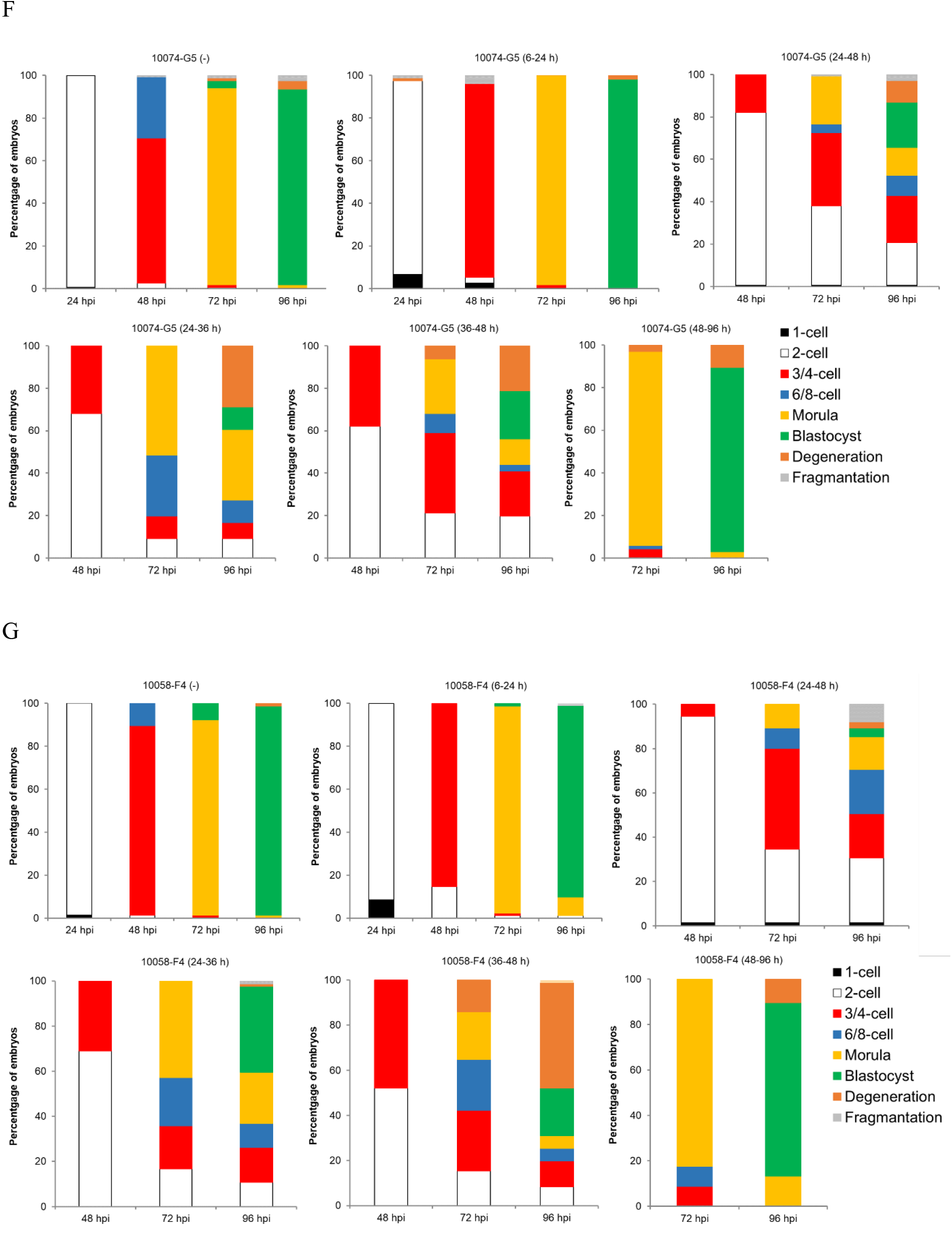
Developmental competence of embryos treated with inhibitor of MYC-MAX heterodimerization. (A) Percentage of embryos by stages of development in 10074-G5 (-) embryos (n=120) or embryos treated with 10074-G5 at the indicated concentrations (2 µM, n=119; 4 µM, n=119, 6 µM, n=119; and 10 µM, n=20) from 6 to 96 h post-insemination (hpi). 10074-G5 (-) indicates the treatment with DMSO, the solvent for 10074-G5, between 6-96 hpi. (B) Representative images of 10074-G5 (-) embryos and embryos treated with 10 µM 10074-G5 from 6 hpi. Embryos were photographed at 24 hpi. Scale bar, 100 μm. (C) Representative images of 10074-G5 (-) embryos or embryos treated with 2, 4 and 6 µM 10074-G5 from 6 hpi. Embryos were photographed at 48 hpi. Scale bar, 100 μm. (D) Percentage of embryos by stages of development in 10058-F4 (-) embryos (n=96) or embryos treated with 10058-F4 at the indicated concentrations (10 µM, n=99; 20 µM, n=100, 30 µM, n=78; and 40 µM, n=80) from 6 to 96 hpi. 10058-F4 (-) indicates the treatment with DMSO, the solvent for 10058-F4, between 6-96 hpi. (E) Representative images of 10058-F4 (-) embryos or embryos treated with 10, 20, 30 and 40 µM 10058-F4 from 6 hpi. Embryos were photographed at 48 hpi. Scale bar, 100 μm. (F) Percentage of embryos by stages of development in 10074-G5 (-) embryos (n=157) or embryos treated with 6 µM 10074-G5 at the indicated periods (6-24 hpi, n=56; 24-48 hpi, n=145, 24-36 hpi, n=66; 36-48 hpi, n=66; and 48-96 hpi n=67). (G) Percentage of embryos by stages of development in 10058-F4 (-) embryos (n=77) or embryos treated with 6 µM 10058-F4 at the indicated periods (6-24 hpi, n=81; 24-48 hpi, n=75, 24-36 hpi, n=84; 36-48 hpi, n=71; and 48-96 hpi n=68).

### MYC-MAX heterodimerization is essential for development at the two-cell stage

To investigate the function of MYC protein in mouse preimplantation development, two structurally distinct small molecule inhibitors, 10074-G5 and 10058-F4 were used in this study. They specifically bind different regions within the helix-loop-helix (HLH) domain of MYC and prevent MYC-MAX heterodimerization. To determine the optimal concentration of 10074-G5 and 10058-F4 with the fewest side effects, embryos were treated with various concentrations of these inhibitors (0, 2, 4, 6 and 10 µM for 10074-G5; 0, 10, 20, 30 and 40 µM for 10058-F4) during 6-96 hpi. Embryos treated with 10 µM 10074-G5 or 30 or 40 µM 10058-F4 were degenerated before cleavage due to cytotoxicity (Figs. 2A, 2B, 2D). Since the developmental rates to the four-cell stage of embryos treated with 6 µM 10074-G5 or 20 µM 10058-F4 were markedly reduced as those treated with *Myc*-ASOs (96.2% for 10074-G5 (-), 16.8% for 10074-G5 (6 µM), 99.0% for 10058-F4 (-) and 27.0% for 10058-F4 (20 µM) at 48 hpi) (Fig. 2A, 2C, 2D), these concentrations were used as the minimum effective concentration in subsequent experiments. To determine the exact time when MYC is required in the development of preimplantation embryos, embryos were treated with the inhibitors for periods of 6-24, 24-48, 24-36, 36-48, or 48-96 hpi. Although inhibitor treatment at 6-24 and 48-96 hpi did not affect embryo development, inhibitor treatments at 24-36, 36-48, and 24-48 hpi reduced the developmental rates to the 4-cell stage, and delayed development was observed in some embryos after transfer to inhibitor-free medium (Fig. 2E and 2F).

### Inhibition of MYC-MAX heterodimerization suppressed ZGA

Gene network analysis predicts that *Myc* is the gene controlling major ZGA^15,16^ We analyzed ChIP-seq data obtained from ESCs^17^ and determined 2,385 MYC direct target genes. Comparing these genes with the 3,025 genes that are elevated during ZGA (ZGA genes)^18^ revealed that 25.9% of ZGA genes are MYC target genes (Fig. 3A). RNA-seq analysis of late two-cell stage embryos treated with 10074-G5 during 6-36 hpi revealed that 82 genes were significantly up-regulated, and 426 genes were significantly down-regulated compared to control embryos (Fig. 3B). Of the down-regulated genes, 152 (35.7%) were direct targets of MYC, and the biological process shown in the gene ontology (GO) analysis of MYC target genes were common in that of down-regulated genes in embryos treated with 10074-G5 (Fig. 3C and 3D). In addition, transcription profiles of embryos treated with 10074-G5 were compared with those of wild-type embryos listed in the public database (DBTMEE)^19^ (Fig. 3E). The results showed that many of the up-regulated genes in 10074-G5 treated embryos were maternal transcripts that were not degraded and genes expressed at minor ZGA phase. In addition, 291 (9.6%) of the ZGA genes were suppressed by 10074-G5 treatment (Fig. 3F). Of the genes that were direct targets of MYC and up-regulated during ZGA, 114 genes were down-regulated due to 10074-G5 treatment (Fig.3G).

**Fig. 3.**
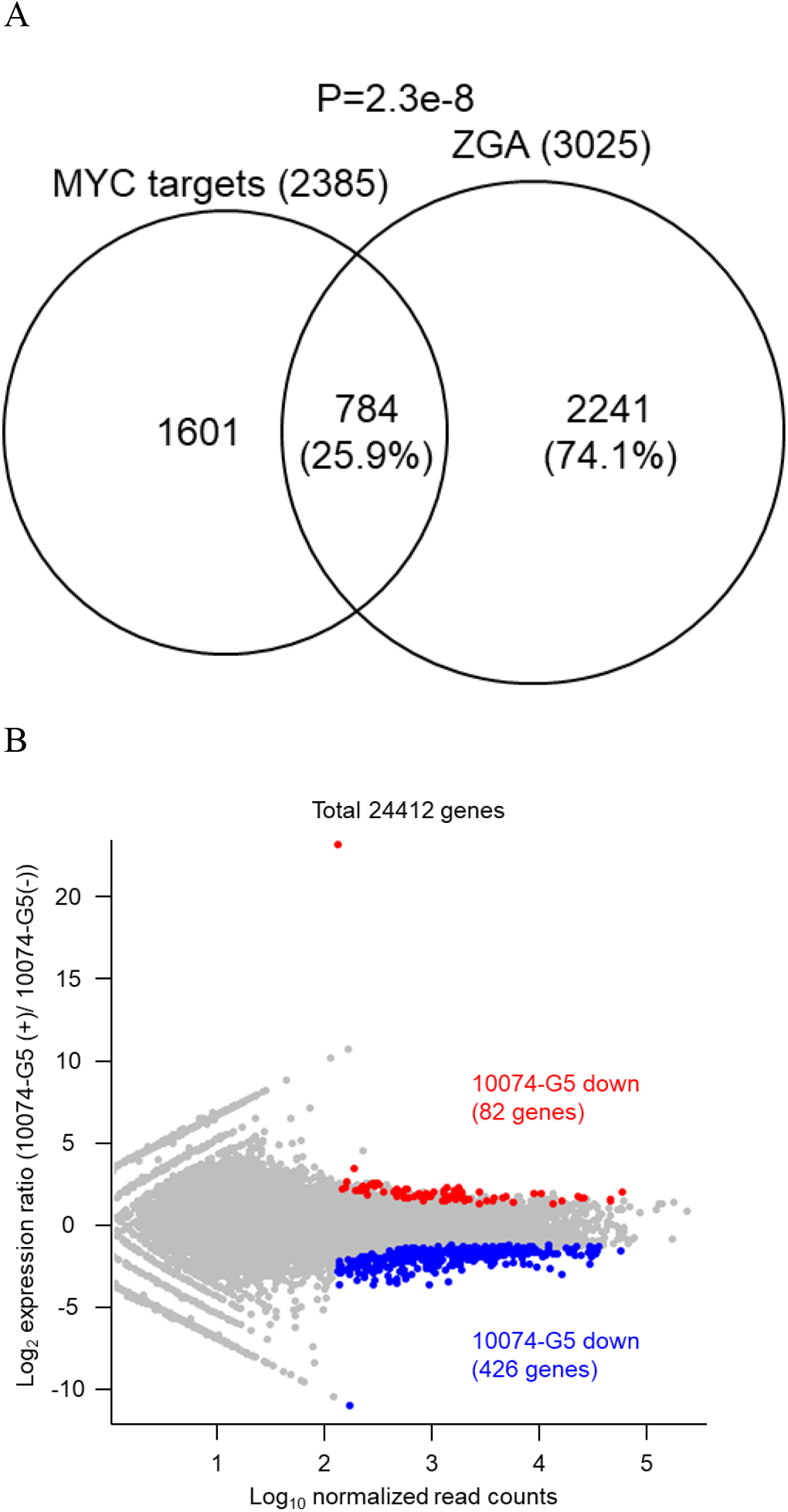
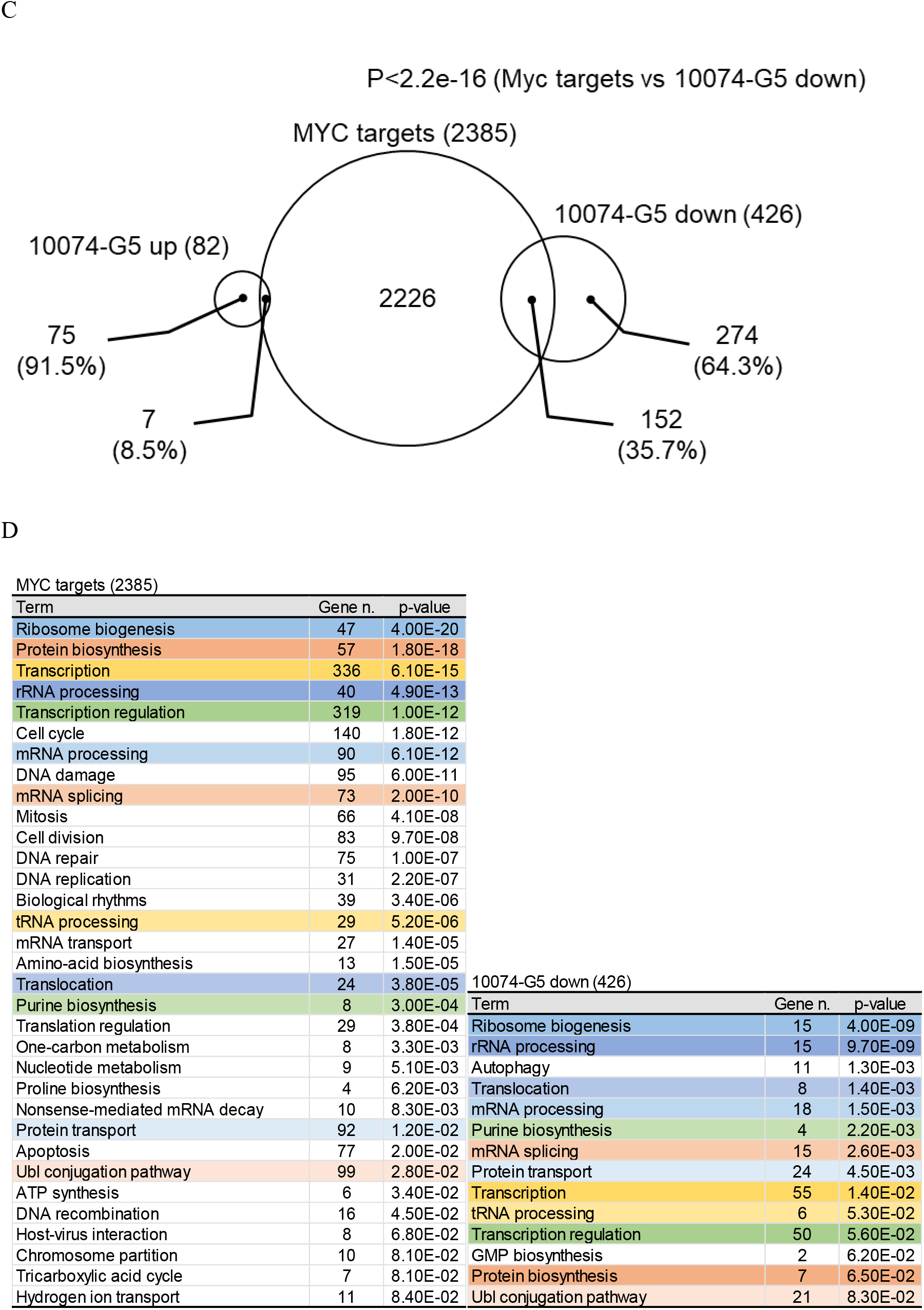
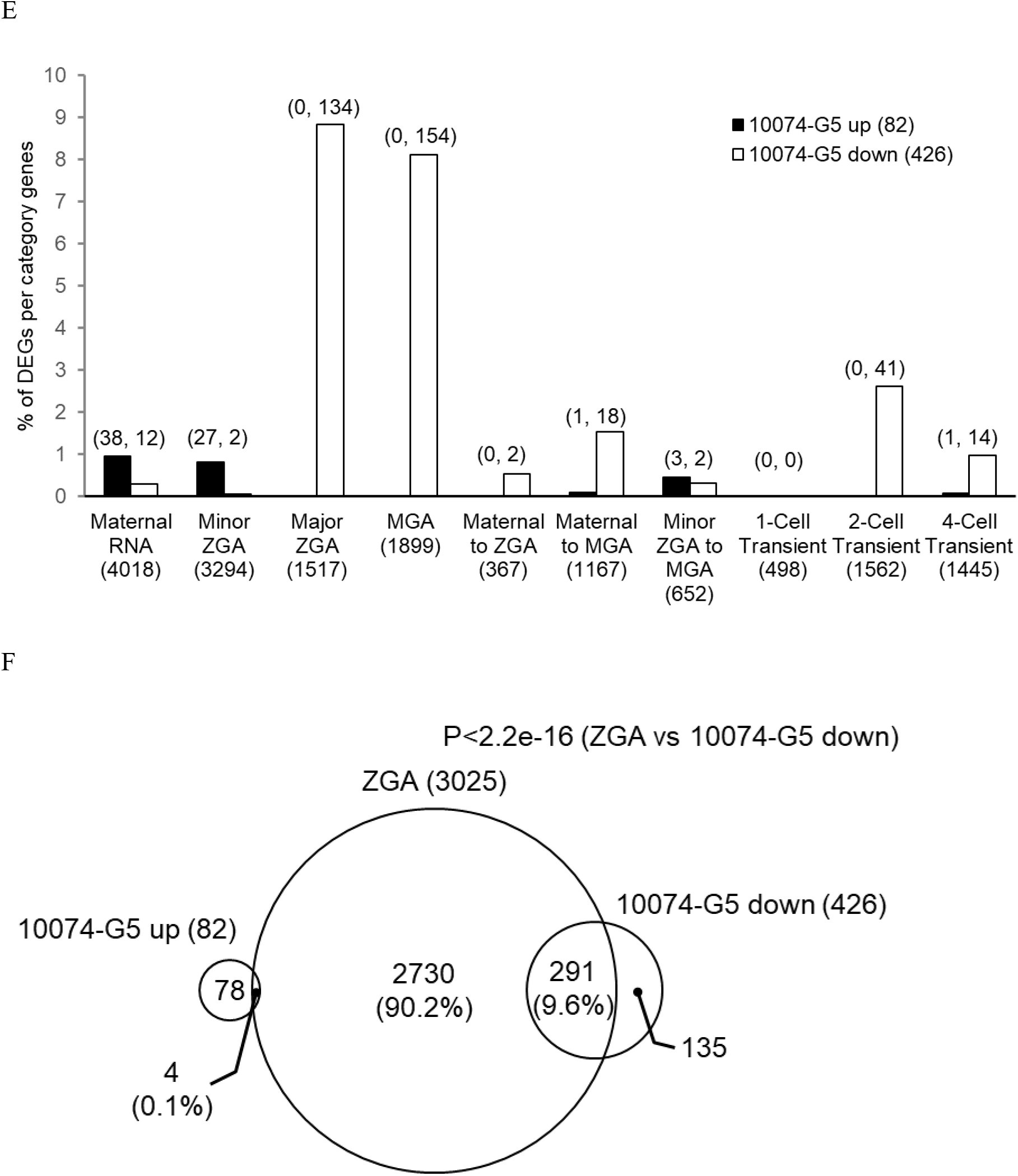
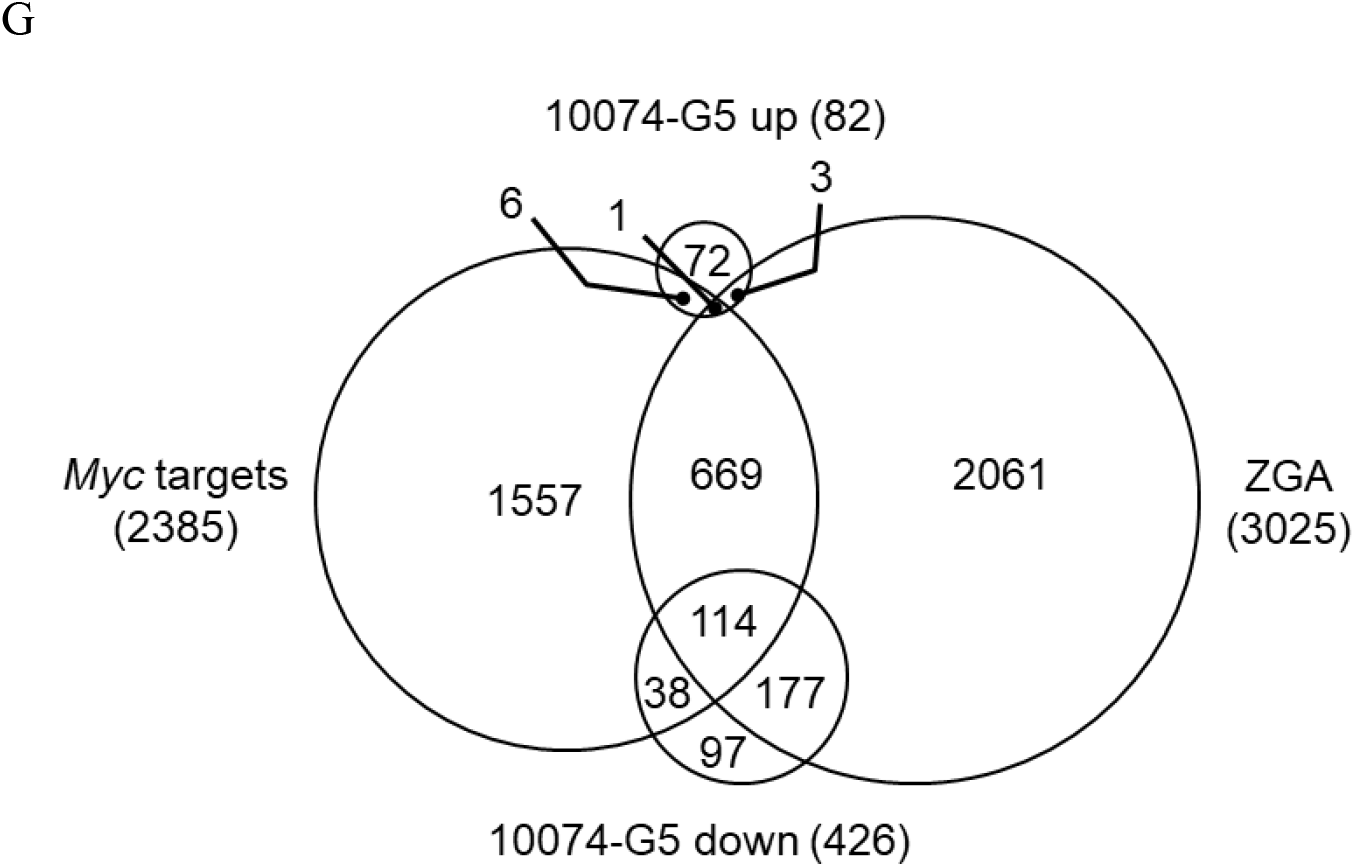
RNA-seq indicates transcriptional aberration by inhibition of MYC-MAX heterodimerization between minor and major ZGA. (A) A Venn diagram between MYC target genes and genes upregulated during ZGA. P values were calculated by Fisher’s exact test. (B) MA plot showing the gene expression ratios of 10074-G5 treated embryos to 10074-G5 (-) embryos and the average gene expression of all embryos. Among 24412 genes, 82 were significantly upregulated (red circles) and 426 were downregulated (blue circles) in 10074-G5 treated embryos. (Padj < 0.05). Differentially expressed genes were defined as those with |log2FC| > 1 and adjusted p value < 0.05 using DESeq2. (C) A Venn diagram between MYC target genes and genes upregulated or downregulated in embryos treated with 10074-G5. P values were calculated by Fisher’s exact test. (D) Significant gene ontology (GO) terms for biological process enriched by the genes with MYC target genes and genes downregulated in embryos treated with 10074-G5. Gene n. represents the numbers of related genes. (E) Percentage of genes upregulated or downregulated in 10074-G5 treated embryos per genes of DBTMEE v2 transcriptome categories. The numbers of genes are indicated above the bars (upregulated genes, downregulated genes). (F) A Venn diagram between genesupregulated in ZGAand genesupregulated or downregulated in embryos treated with 10074-G5. P values were calculated by Fisher’s exact test. (G) A Venn diagram between MYC target genes and genes upregulated in ZGA and genes upregulated or downregulated in embryos treated with 10074-G5.

## Discussion

This study demonstrates that the transcription factor MYC is essential for early mouse embryogenesis by using ASOs and inhibitors. In this study, we found that developmental arrest occurs at the two-cell stage when treated with ASOs or MYC inhibitors from the one-cell stage. The functions of *Mycn* and *Mycl* are redundant with those of *Myc*, and embryos in which *Myc* on a chromosome is replaced with *Mycn* develop normally^20^. Although the inhibitors used in this study have been shown to be effective against MYCN and MYCL^21,22^, developmental arrest occurred at the two-cell stage even with ASOs targeting only *Myc*, suggesting that only *Myc* is required for early development, and this idea is also valid since RNA-seq data analysis indicates that *Mycn* and *Mycl* expression is low in early development. Although Paria et al. showed that *Myc*-targeted ASO-treatment from the two-cell stage resulted in embryonic arrest at the eight-cell/morula stage^11^, some embryos were similarly arrested at the eight-cell/morula stage (10074-G5: 80.3% and 10058-F4: 33.3%) when the two-cell embryos were treated with MYC inhibitors at a limited-time (24-36 hpi) in this study. This suggests that major ZGA at the two-cell stage induced by MYC was partially inhibited in embryos arrested at the eight-cell/morula stage by ASO treatment in the previous study and by the time-limited inhibitor treatment in the present study. More recently, it was reported that treatment of mouse one-cell embryos with 10058-F4, the same inhibitor of MYC that we used, resulted in developmental arrest at the one-cell stage^13^. Since it has been reported that inhibition of minor ZGA alone does not impair development from the one-cell stage to the two-cell stage, but impairs development after the two-cell stage^23^, we hypothesized that inhibition of the transcription factor including *Myc* would not result in one-cell stage arrest.

In our study, the optimal inhibitor concentration was the concentration at which development stops at the two-cell stage, similar to the results with ASOs, but when treated with a higher concentration of 10058-F4, as was the case in their study, developmental arrest or degeneration also occurred at the one-cell stage, suggesting that the effect of the higher inhibitor concentration on development may be an artifact. In our research, embryos developed normally into blastocysts when treated with inhibitors at the one-to two-cell stage (6-24 hpi) or after the four-cell stage (48-96 hpi), also suggesting that the inhibitors specifically inhibited MYC without cytotoxic effects.

RNA-seq analysis of 10074-G5-treated embryos during 6-36 hpi (from just after fertilization to the late-2-cell stage) showed that only two genes classified as minor ZGA were down-regulated, while many (289) genes classified as major ZGA and MGA were down-regulated. This indicates that MYC is required for induction of major ZGA, as pointed out in previous studies using gene network analysis^15,16^, and that the transcriptional aberration caused by MYC inhibition occurred between minor ZGA and major ZGA. This result is supported by a report that MYC proteins co-localize with nuclear speckles in granular form in the nucleus from the two-cell stage^10^, suggesting that MYC localizes to transcriptionally active regions around nuclear speckles and promotes transcription of the embryonic genome. On the other hand, an increase in the number of genes classified as maternal RNAs was observed by RNA-seq analysis, but this result was attributed to the lack of degradation of maternal transcripts. The number of such genes was only 38, indicating that the effect of MYC inhibition on the degradation of maternal transcripts was relatively small. GO analysis of MYC target genes and genes whose expression was downregulated by MYC inhibition showed that both contain many genes involved in pathways such as ribosome biosynthesis, protein biosynthesis and transcription. These results suggest that the downstream genes of MYC dominate central dogma in major ZGA, and developmental arrest occurred because these important pathways were disrupted.

Of the 2385 direct MYC target genes obtained from chip-seq data in ES cells, only 152 genes (6.37%) were down-regulated at the two-cell stage by MYC inhibition. The genes whose expression was not decreased may be due to differences in MYC target genes in ES cells and early embryos. Indeed, among the top 500 genes that were highly expressed in DMSO-treated control embryos, 167 (33.4%) were MYC target genes, and 44 (26.3%), a higher proportion than overall MYC target genes in ES cells, were down-regulated by MYC inhibition. In addition, based on the report that when the relative amount of MYC is low, MYC binds only to high-affinity targets, and when the relative amount is high, MYC also binds to low-affinity targets^23,24^, it is possible that the difference in the relative amount of MYC between ES cells and early embryos was the reason for the low number of genes whose expression was decreased by MYC inhibition at the two-cell stage.

*Myc* is one of the proto-oncogenes commonly expressed in various types of cancer cells, and the feature that both cancer cells and early embryos lack cell cycle checkpoints and proliferate rapidly may be derived from *Myc*^25^. *Myc* is also one of the genes selected for establishment of iPS cells because it maintains the pluripotency of ES cells, and expression of four genes including *Myc* can reprogram fibroblasts into pluripotent stem cells^26^. On the other hand, it has been shown that 2-cell-like cells are induced from ES cells when *Myc* is knocked out, suggesting that *Myc* prevents the transition from the pluripotent state to the totipotent state in ES cells^27^.

The present study demonstrates that *Myc* is required at the 2-cell stage, the time when the pluripotent state is lost and the transition to the pluripotent state is initiated, suggesting that *Myc* also affects the differentiation potential of the early embryo. Our understanding of the “essential” role of *Myc* in early embryogenesis will accelerate our understanding of cancer cells and induced pluripotent cells.

## Materials and Methods

### Collection of oocytes, in vitro fertilization, and embryo culture

8–12-week-old ICR mice (Japan SLC, Shizuoka, Japan) were superovulated by injection with 7.5 IU equine chorionic gonadotropin (eCG; ASUKA Animal Health, Tokyo, Japan) followed 46-48 h later by 7.5 IU of human chorionic gonadotropin (hCG; ASUKA Animal Health). Cumulus-oocyte complexes (COCs) were collected from the ampullae of the excised oviducts 14 h after the hCG injection. COCs were placed in a 100-μl droplet of human tubal fluid (HTF) medium supplemented with 4 mg/ml BSA (Sigma-Aldrich, St. Louis, MO, USA)^28^. Spermatozoa were collected from the cauda epididymis of 12-to 18-week-old ICR male mice (Japan SLC) and cultured for at least 1 h in a 100 -μl droplet of HTF. After preincubation, sperm suspension was added to fertilization droplets at a final concentration of 1 × 10^6^ cells/ml. At 6 hpi, morphologically normal fertilized embryos were collected and cultured in potassium simplex optimized medium (KSOM) supplemented with amino acids^29^ and 1 mg/ml BSA under paraffin oil (Nacalai Tesque, Kyoto, Japan). All incubations were performed at 37°C under 5% CO_2_.

### RNA extraction and reverse transcription-quantitative PCR (RT-qPCR)

Total RNA extraction and cDNA synthesis from oocytes or cumulus cells were performed using the superPrepTM Cell Lysis & RT Kit for qPCR (TOYOBO, Osaka, Japan). Synthesized cDNA was mixed with specific primers and KOD SYBR qPCR Mix (TOYOBO), followed by RT-qPCR amplification. The protocols for RT-qPCR and establishment of transcript levels were performed as previously described^30^, and *Gapdh* or *H2afz* was used as an internal control. The relative gene expression was calculated using the 2−ΔΔCt method^31^. Primer sequences used for RT-qPCR were as follows: *Gapdh*, 5 ′-GTGTTCCTACCCCCAATGTG-3 ′(forward) and 5 ′-TGTCATCATACTTGGCAGGTTTC-3 (reverse); *H2afz*, 5 ′-TCCAGTGGACTGTATCTCTGTGA-3 ′(forward) and 5 ′-GACTCGAATGCAGAAATTTGG-3 ′(reverse); *Myc*, 5 ′-CGTTGGAAACCCCGCAGA-3 ′(forward) and 5 ′-TCCAGATATCCTCACTGGGCG-3 ′(reverse).

### Immunocytochemistry and fluorescence analysis

To detect the localization of MYC, embryos were fixed with 4% paraformaldehyde in phosphate-buffered saline (PBS) for 20 min at 28°C. Then embryos were treated with 0.5% Triton X-100 (Sigma-Aldrich) in PBS for 40 min at 28°C. Embryos were blocked in PBS containing 1.5% BSA, 0.2% sodium azide, and 0.02% Tween20 (antibody dilution buffer) for 1 h at 28°C, followed by overnight incubation at 4°C with rabbit anti-MYC (1:20,000) (ab32072; Abcam Ltd, Cambridge, UK) antibody in antibody dilution buffer. Subsequently, the samples were washed in antibody dilution buffer and incubated with Alexa Fluor 594 conjugated goat anti-rabbit IgG secondary antibody (1:500; Thermo Fisher Scientific, Waltham, MA, USA) for 1 h at 28°C. After washing in antibody dilution buffer, embryos were stained with antibody dilution buffer containing 10 μg/ml Hoechst 33342 (Sigma-Aldrich) for 20 min at 28°C. Stained embryos were mounted on glass slides, and signals were observed using a fluorescence microscope (IX73; Olympus, Tokyo, Japan).

### Microinjection of antisense oligonucleotides

ASOs were purchased from Qiagen (Hilden, Germany). *Myc*-ASO-1, LG00828024-DDA; *Myc*-ASO-2, LG00828026-DDA; and NT-ASO LG00000002-DDA. Approximately 3–5 pl of 10 µM ASOs were microinjected into the cytoplasm of zygotes at 3 hpi.

### Inhibitor treatment

10074-G5 (Cayman Chemical Company, Ann Arbor, MI, USA) and 10058-F4 (Cayman Chemical Company) are MYC inhibitors that bind to and distort the bHLH-ZIP domain of MYC, thereby inhibiting MYC/MAX heterodimer formation and inhibiting its transcriptional activity^32^. These two inhibitors are structurally unrelated and their cognate binding sites on MYC are distinct from each other. Embryos were cultured in KSOM containing 2, 4, 6, or 10 µM 10074-G5 or 10, 20, 30, or 40 µM 10058-F4 between 6-24, 6-96, 24-36, 24-48, 36-48, or 48-96 hpi. Control embryos were cultured in inhibitor-free medium that contained the appropriate amount of dimethyl sulfoxide (DMSO), the solvent used to prepare the stock solution of inhibitors.

### Publicly available data

The MYC ChIP-seq data were downloaded from ref.^17^ with GEO accession number GSM1171648. Raw reads were aligned to the mm10 genome using Bowtie 2 (v2.4.5)^33^. Removing mapping duplicates and handling of sam and bam files were performed by SAMtools (v1.10)^34^. The peaks were called using MACS2 (v2.2.7.1)^35^.

### RNA-seq and data processing

Two-cell stage embryos (n=10) in the control and inhibitor-treated groups were collected for RNA-seq library construction at 36 hpi. Two biological replicates were obtained from each group. Polyadenylated RNA-seq libraries were prepared by using SMART-Seq v4 PLUS Kit (Clontech, Mountain View, CA, USA). Indexed RNA-seq libraries were sequenced using Illumina HiSeqX sequencer (paired end, 150 bp). The resulting sequence reads were mapped to the mm10 mouse genome using STAR aligner software (v2.7.10.a) and only uniquely mapped reads were used for subsequent processing^36^. Gene expression levels were calculated using RSEM (v1.3.3)^37^ and differentially expressed genes were analyzed by the DESeq2 package (version 1.38.3)^38^. Differentially expressed genes were defined as those with |log2FC| > 1 and adjusted p value < 0.05. Gene ontology (GO) analysis was performed using the DAVID tool^39,40^.

### Statistical analyses

Developmental rates were analyzed using the chi-squared test with Holm’s adjustment. For gene expression analysis, RT-qPCR data were analyzed using one-way analysis of variance (ANOVA) followed by the Tukey-Kramer test. *P* values <0.05 were considered statistically significant.

### Ethical approval for the use of animals

All experimental procedures were approved by the Animal Research Committee of Kyoto University (Permit no. R3-17) and performed in accordance with the committee’s guidelines.

## Data availability statement

The original contributions presented in the study are publicly available. This data can be found here: https://www.ncbi.nlm.nih.gov/geo/query/acc.cgi?acc=GSE233897

## Acknowledgements

This work was supported by Grant-in-Aid for Scientific Research (no. 19H03136 to NM) and Grant-in-Aid for JSPS Fellows (no. 21J21840 to TY) from the Japan Society for the Promotion of Science.

## Author contributions

TY, HW, SH, SI, and NM conceived the study. TY, HW, and HS performed the experiments and analyzed the data. TY, SH, SI, and NM wrote the manuscript. All authors discussed the re sults and approved the manuscript.

## Reference

1. Blackwood, E. M. & Eisenman, R. N. Max: A Helix-Loop-Helix Zipper Protein That Forms a Sequence-Specific DNA-Binding Complex with Myc. Science (1979) 251, 1211–1217 (1991).

2. Rahl, P. B. et al. c-Myc Regulates Transcriptional Pause Release. Cell 141, 432–445 (2010).

3. Zhang, K. & Smith, G. W. Maternal control of early embryogenesis in mammals. Reprod Fertil Dev 27, 880–896 (2015).

4. Tadros, W. & Lipshitz, H. D. The maternal-to-zygotic transition: a play in two acts. Development 136, 3033–3042 (2009).

5. Li, L., Zheng, P. & Dean, J. Maternal control of early mouse development. Development 137, 859–870 (2010).

6. Aoki, F., Worrad, D. M. & Schultz, R. M. Regulation of Transcriptional Activity during the First and Second Cell Cycles in the Preimplantation Mouse Embryo. Dev Biol 181, 296–307 (1997).

7. Minami, N., Suzuki, T. & Tsukamoto, S. Zygotic Gene Activation and Maternal Factors in Mammals. Journal of Reproduction and Development 53, 707–715 (2007).

8. Schulz, K. N. & Harrison, M. M. Mechanisms regulating zygotic genome activation. Nature Reviews Genetics 2018 20:4 20, 221–234 (2018).

9. Jukam, D., Shariati, S. A. M. & Skotheim, J. M. Zygotic Genome Activation in Vertebrates. Dev Cell 42, 316–332 (2017).

10. Suzuki, T., Abe, K. I., Inoue, A. & Aoki, F. Expression of c-MYC in Nuclear Speckles During Mouse Oocyte Growth and Preimplantation Development. Journal of Reproduction and Development 55, 491–495 (2009).

11. Paria, B. C., Dey, S. K. & Andrews, G. K. Antisense c-myc effects on preimplantation mouse embryo development. Proc Natl Acad Sci U S A 89, 10051 (1992).

12. Davis, A. C., Wims, M., Spotts, G. D., Hann, S. R. & Bradley, A. A null c-myc mutation causes lethality before 10.5 days of gestation in homozygotes and reduced fertility in heterozygous female mice. Genes Dev 7, 671–682 (1993).

13. Asami, M. et al. A program of successive gene expression in mouse one-cell embryos. Cell Rep 42, 112023 (2023).

14. Kinisu, M. et al. Klf5 establishes bi-potential cell fate by dual regulation of ICM and TE specification genes. Cell Rep 37, 109982 (2021).

15. Aoki, F. Zygotic gene activation in mice: profile and regulation. J Reprod Dev 68, 79 (2022).

16. Zeng, F. & Schultz, R. M. RNA transcript profiling during zygotic gene activation in the preimplantation mouse embryo. Dev Biol 283, 40–57 (2005).

17. Krepelova, A., Neri, F., Maldotti, M., Rapelli, S. & Oliviero, S. Myc and Max Genome-Wide Binding Sites Analysis Links the Myc Regulatory Network with the Polycomb and the Core Pluripotency Networks in Mouse Embryonic Stem Cells. PLoS One 9, 88933 (2014).

18. Arand, J. et al. Tet enzymes are essential for early embryogenesis and completion of embryonic genome activation. EMBO Rep 23, (2022).

19. Park, S. J., Shirahige, K., Ohsugi, M. & Nakai, K. DBTMEE: a database of transcriptome in mouse early embryos. Nucleic Acids Res 43, D771 (2015).

20. Malynn, B. A. et al. N-myc can functionally replace c-myc in murine development, cellular growth, and differentiation. Genes Dev 14, 1390 (2000).

21. Müller, I. et al. Targeting of the MYCN Protein with Small Molecule c-MYC Inhibitors. PLoS One 9, e97285 (2014).

22. Fletcher, S. & Prochownik, E. V. Small-molecule inhibitors of the Myc oncoprotein. Biochimica et Biophysica Acta (BBA) - Gene Regulatory Mechanisms 1849, 525–543 (2015).

23. Lin, C. Y. et al. Transcriptional Amplification in Tumor Cells with Elevated c-Myc. Cell 151, 56 (2012).

24. Fernandez, P. C. et al. Genomic targets of the human c-Myc protein. Genes Dev 17, 1115 (2003).

25. Dang, C. V. MYC on the Path to Cancer. Cell 149, 22 (2012).

26. Takahashi, K. & Yamanaka, S. Induction of Pluripotent Stem Cells from Mouse Embryonic and Adult Fibroblast Cultures by Defined Factors. Cell 126, 663–676 (2006).

27. Fu, X., Wu, X., Djekidel, M. N. & Zhang, Y. Myc and Dnmt1 impede the pluripotent to totipotent state transition in embryonic stem cells. Nat Cell Biol 21, 835 (2019).

28. Minami, N., Sasaki, K., Aizawa, A., Miyamoto, M. & Imai, H. Analysis of Gene Expression in Mouse 2-Cell Embryos Using Fluorescein Differential Display: Comparison of Culture Environments. Biol Reprod 64, 30–35 (2001).

29. Ho, Y., Wigglesworth, K., Eppig, J. J. & Schultz, R. M. Preimplantation development of mouse embryos in KSOM: Augmentation by amino acids and analysis of gene expression. Mol Reprod Dev 41, 232–238 (1995).

30. Shikata, D., Yamamoto, T., Honda, S., Ikeda, S. & Minami, N. H4K20 monomethylation inhibition causes loss of genomic integrity in mouse preimplantation embryos. J Reprod Dev 66, 411–419 (2020).

31. Livak, K. J. & Schmittgen, T. D. Analysis of relative gene expression data using real-time quantitative PCR and the 2-ΔΔCT method. Methods 25, 402–408 (2001).

32. Yin, X., Giap, C., Lazo, J. S. & Prochownik, E. V. Low molecular weight inhibitors of Myc - Max interaction and function. Oncogene 22, 6151–6159 (2003).

33. Langmead, B. & Salzberg, S. L. Fast gapped-read alignment with Bowtie 2. Nat Methods 9, 357–359 (2012).

34. Li, H. et al. The Sequence Alignment/Map format and SAMtools. Bioinformatics 25, 2078–2079 (2009).

35. Zhang, Y. et al. Model-based analysis of ChIP-Seq (MACS). Genome Biol 9, 1–9 (2008).

36. Dobin, A. et al. STAR: ultrafast universal RNA-seq aligner. Bioinformatics 29, 15 (2013).

37. Li, B. & Dewey, C. N. RSEM: accurate transcript quantification from RNA-Seq data with or without a reference genome. BMC Bioinformatics 12, 323 (2011).

38. Love, M. I., Huber, W. & Anders, S. Moderated estimation of fold change and dispersion for RNA-seq data with DESeq2. Genome Biol 15, 550 (2014).

39. Huang, D. W., Sherman, B. T. & Lempicki, R. A. Systematic and integrative analysis of large gene lists using DAVID bioinformatics resources. Nat Protoc 4, 44–57 (2009).

40. Huang, D. W., Sherman, B. T. & Lempicki, R. A. Bioinformatics enrichment tools: paths toward the comprehensive functional analysis of large gene lists. Nucleic Acids Res 37, 1 (2009).

